# Hedgehog Pathway Activity Downstream of Smoothened is regulated specifically by Basal Ciliary PKA

**DOI:** 10.1101/2024.12.30.630707

**Authors:** Hongyu Zhang, Shujing Chen, Zhuoya Huang, Jin Ben, Guangxin Chen, Changxin Wu, Haibo Xie, Philip Ingham, Zhonghua Zhao

## Abstract

Effectors of the vertebrate Hedgehog (HH) signaling pathway are organized through primary cilia (PC) that grow and retract in lockstep with the cell cycle in response to extracellular signals. Protein kinase A (PKA), a kinase with ubiquitous distribution in most cells, functions as a specific negative regulator of the HH pathway. Its functional specificity in the HH pathway has been suggested to be controlled by cAMP in the PC. However, the regulation of PKA and its functions in PC remain unclear, in part due to the lack of observation of PKA localization in PC during HH resting state as well as conflicting reports of the dynamic changes of cAMP in cilia and HH pathway activity. To address this issue, we have developed a ciliary-localized FRET-based A-kinase activity probe (Nphp3N-AKAR2-CR) as an improved biosensor for monitoring real-time PKA activity in the PC of both cultured cells and living zebrafish embryos. Although the PKA catalytic subunit (PKA-C) was not observed in PC, basal PKA activity in cells could be detected with this probe. In addition, we have found that only ciliary-targeted PKA and not cytosolic PKA, can modulate the HH pathway, even when the integrity of the PC is disrupted. Notably, ciliary PKA activity was barely changed either by inhibition or activation of the HH pathway at the level of Smoothened (SMO), the obligate HH signal transducer. Moreover, we found that even low concentration of the adenylyl cyclase agonist forskolin (FSK) can efficiently inhibit the HH pathway in the presence of the constitutively active variant SMOA1, suggesting that the activation of the HH pathway by SMO may not be due solely to direct regulation of PKA activity in the PC.

## Introduction

The HH signaling pathway is conserved from cnidaria to vertebrates, playing critical roles in embryonic development and adult tissue and metabolic homeostasis (Ingham, 2022). Abnormal HH pathway activity results in multiple developmental defects and is also implicated in the onset and progression of various malignant cancers (Barakat et al., 2010; Scales and Sauvage, 2009; Taipale and Beachy, 2001). The mechanism of HH signal transduction has been extensively studied in vertebrate systems, including mice, zebrafish and cultured cells, as well as invertebrates, especially *Drosophila,* in which it was first characterized (Ingham, 2022). Most of the core components of the pathway are shared between phyla and have highly similar mechanisms controlling the transduction of the HH signal. However, the process is more complex in vertebrates, notably through its dependence on the primary cilium (PC), a highly specialized subcellular compartment.

A major event in PC regulation of HH pathway activity is the divergent phosphorylation of the GLI proteins, generating either the full-length active form or truncated repressor form (Rimkus et al., 2016; Dilower et al., 2023). Protein Kinase A (PKA), one of the key negative regulators of the HH pathway, is considered to be the critical kinase for phosphorylating GLI and promoting its cleavage to a repressor form, thereby inhibiting HH pathway target gene activation (Concordet et al., 1996; Johnson et al., 1995; Lepage et al., 1995; Li et al., 1995; Hammerschmidt et al., 1996; Robbins et al., 1997; Chen et al., 1998; Wang et al., 1999; Zhang et al., 2015).

The PKA holoenzyme is a heterotetramer composed of two catalytic subunits (PKA-C) and two regulatory subunits (PKA-R), that are ubiquitously expressed and involved in many intracellular signaling pathways mediating numerous physiological processes, including cell proliferation, metabolism, memory formation and muscle function (Rosenberg et al., 2002; Bollen et al., 2014; Torella et al., 2009). The localization of PKA to specific subcellular compartments is crucial for its diverse signaling functions (Scott and Pawson, 2009). The activity of PKA is primarily controlled by cyclic adenosine monophosphate (cAMP), which binds directly to the PKA-R subunit and thereby releases active PKA-C (Torres-Quesada et al., 2017). Generally, cAMP level is dependent upon Adenylyl Cyclases (ACs) that are either activated or inhibited by the α_s_ or α_i_ G-protein subunits, respectively, in response to G-Protein-Coupled Receptor (GPCR) activation. Furthermore, the heat-stable protein kinase inhibitor (PKI) proteins can also modulate PKA activity through direct interaction with the catalytic subunit of PKA-C (Dalton and Dewey, 2006). In the HH pathway, Smoothened (SMO), an F-family GPCR, has been hypothesized to inhibit PKA activity, either directly by sequestering the PKA-C subunit via a pseudosubstrate domain in its intracellular C-terminal domain (Happ et al., 2022; Walker et al., 2024) or indirectly, by activating Gα_i_ (Ogden et al., 2008; DeCamp et al., 2000; Shen et al., 2013). GPR161, a typical ciliary GPCR, has also been implicated in modulating HH pathway activity, by increasing PKA activity through Gα_s_ in the ventral neural tube, as well as in chondrocytes and other ciliated cells (Mukhopadhyay et al., 2013; Hwang et al., 2018). Both GPCRs are localized to the PC and show opposing ciliary accumulation in response to HH signaling, consistent with the regulation of HH pathway activity by PKA occurring within the PC. In line with this, the PKA-R subunit has been directly visualized in the PC (Mick et al., 2015). By contrast, although ciliary localization of the PKA-C subunit has been inferred from proteomic analysis (Mick et al., 2015), it has only been visualised in the PC following HH activation, coincident with the translocation of SMO (Walker et al., 2024). Such distribution dynamics seem paradoxical if HH pathway activity is supposed to be suppressed by PKA within the PC. Moreover, the prevailing model proposing that PC maintain relatively higher cAMP levels compared to the cytoplasm in the HH-OFF state, followed by a reduction in cAMP upon HH pathway activation, remains controversial, further obscuring the association between ciliary PKA and HH pathway activity (Moore et al., 2016; Jiang et al., 2019).

Previous studies have used the 5HT_6_-AKAR4 to assay PKA activity in the PC (Moore et al., 2016). By targeting the FRET-based A-kinase activity probe (AKAR2) to the PC using the N-terminal region of zebrafish Nephrocystin-3 (Nphp3) (Zhang et al, 2022), we have established an improved biosensor that facilitates the real-time dynamic monitoring of PKA in the PC of both cultured cells and living embryos. The sensor can detect changes in PKA activity in the PC in response to forskolin (FSK) treatment as well as to the expression of the dominant negative PKA-R variant, confirming a basal level of PKA activity in the PC. Furthermore, by utilizing the Nphp3N to deliver different PKA variants specifically to either the PC or the cytoplasm in both cells and embryos, we have found that only ciliary-targeted PKA specifically regulates the HH pathway, even in the absence of the PC. Unexpectedly, however, perturbation of the HH pathway at the level of SMO did not change PKA activity within the PC. Moreover, constitutive activation of SMO was not sufficient to overcome the effects of increased activation of PKA mediated by a range of FSK concentrations, contrary to expectation if SMO inhibits PKA through directly sequestering it in PC. Taken together, our findings suggest that a basal level of PKA localized in the PC and not the cytoplasm, specifically regulates HH signal transduction in cells.

## Results

### Construction of a biosensor that accurately reports cilia-specific PKA activity

The AKAR2-CR biosensor is a FRET-based Protein kinase A (PKA) activity probe developed to monitor PKA activity in cells (Ni et al., 2006; Allen and Zhang, 2006). It is composed of a fusion protein consisting of a phospho-peptide binding domain, a Forkhead-associated domain (FHA1), and a consensus region of PKA substrates sandwiched between a donor fluorophore (Clover) and an acceptor fluorophore (mRuby2). In the absence of PKA activity, AKAR2-CR maintains a straight structure such that the Clover and the mRuby2 fluorophores are separated, leading to low FRET efficiency. When the central substrate is phosphorylated by PKA-C, the AKAR2-CR folds, bringing the two fluorophores into close enough proximity for FRET to occur efficiently. Thus, AKAR2-CR can accurately report PKA activity within cells (Fig. 1B).

**Figure 1.**
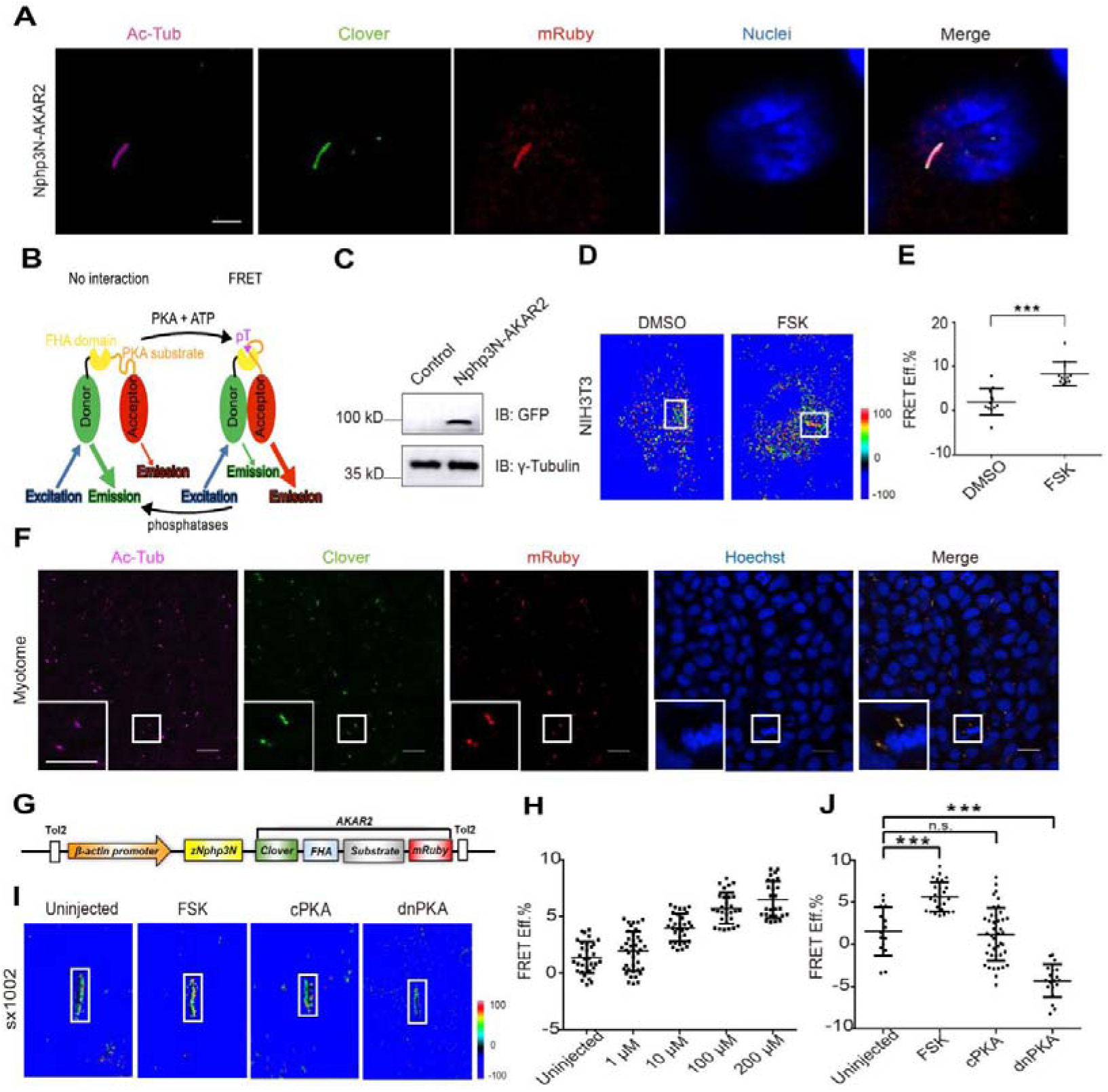
The Nphp3N-AKAR2-CR biosensor can accurately and efficiently report PKA activity in the PC. (A) Specifically ciliary localization of Nphp3N-AKAR2-CR as indicated by colocalization of Clover in green, mRuby in red with cilium stained by acetylated tubulin in purple. The nuclei were labelled in blue by Hoechst. Scale bars, 5 μm. (B) Schematic model of the AKAR2-CR illustrating the PKA activity. Upon phosphorylation by PKA, the substrate region binds FHA domain, thereby facilitating the assembly of the donor and acceptor and leading to FRET. The phosphorylated reporter increased the ratio of acceptor:donor emission and reduced the lifetime of donor fluorescence. (C) The expression of Nphp3N-AKAR2-CR was detected by immunoblot using GFP antibody against AKAR2-CR. The γ-tubulin was used as loading control. (D) The FRET ratio image of ciliary AKAR2-CR in NIH3T3 cells treated with DMSO and 10 μM forskolin (FSK), respectively. (E) Statistical analysis of the ciliary FRET efficiency in NIH3T3 cells in response to DMSO and FSK, respectively (n_DMSO_=12 cilia, n_FSK_=12 cilia). One-way ANOVA was performed, and *** indicates ‘significant’ with *P*<0.001. (F) The stable transgenic line embryos indicated by Clover in green and mRuby in red were efficiently localized in the primary cilia of cells in the myotome. Cilia were labeled by Ac-Tub in magenta. Each small frame on the lower right corner of the main frame denotes the enlarged site as encircled by white square. The nuclei were labelled by Hoechst in blue. Scale bar, 5 μm. (G) Schematic diagram of the *Nphp3N-AKAR2-CR* vector used for generating the transgenic ciliary PKA reporter zebrafish. (H) Statistical analysis of the ciliary FRET efficiency in *sx1002* treated with gradient-diluted FSK (n_untreated_=32 cilia, n_1μm_=40 cilia, n_10μm_=34 cilia, n_100μm_ =30 cilia, n_200μm_ =27 cilia). (I) The FRET ratio image of ciliary AKAR2-CR in *sx1002* injected with FSK, cPKA and dnPKA, respectively. (J) Statistical analysis of the ciliary FRET efficiency in *sx1002* treated with FSK, cPKA and dnPKA (n_uninjected_=18 cilia, n_FSK_=30 cilia, n_cPKA_=40 cilia, n_dnPKA_=24 cilia). One-way ANOVA was performed, and the ns indicates ‘not significant’ with P>0.05, while *** indicates ‘significant’ with *P*<0.001.

To develop a specific PKA biosensor of the PC, we fused, Nphp3N, which has been shown to be a specific and efficient ciliary targeting peptide (CTP), to the N terminus of AKAR2-CR. When introduced into NIH3T3 cells, the Nphp3N-AKAR2-CR fusion protein was normally expressed and specifically localized to the PC (Fig. 1A and 1C), as indicated by the overlap of Clover and mRuby2 signals with acetylated tubulin, a cilia marker (Fig. 1A). Treatment of transfected cells with FSK, led to a ∼9% increase in FRET efficiency in the PC (Fig. 1D and 1E), demonstrating that the Nphp3N-AKAR2-CR biosensor can efficiently report PKA activity increase in the PC of NIH3T3 cells.

The ease of visualization of early embryonic stages in the zebrafish together with robust readouts for cilia and HH pathway activity (Barresi et al., 2000; Wolff et al., 2003; Leventea et al., 2016; Jaffe et al., 2010), makes it a good model for investigating the role of PKA in HH signaling in vivo. To expand the ciliary PKA biosensor to organism level, the expression and PKA biosensor activity of Nphp3N-AKAR2-CR has been examined. Transient expression of the Nphp3N-AKAR2-CR in zebrafish embryos showed that it localizes specifically in the PC as in NIH3T3 cells (Fig. S1A). However, the levels of expression varied between cells. To overcome this variability, we generated a stable transgenic line in which Nphp3N-AKAR2-CR is ubiquitously expressed throughout the embryo, driven by the β*-actin* promoter (Fig. 1G) (Higashijima et al., 1997), and designated it as Tg (β*-actin*::*nphp3N-AKAR2-CR*)^sx1002^ (*sx1002* hereafter). The *sx1002* fish develop normally and are fertile. Nearly all the PC of a variety of cells analyzed in *sx1002* embryos exhibited Clover and mRuby2 signal simultaneously (Fig. 1F), suggesting that the Nphp3N-AKAR2-CR fusion protein maintained stable and persistent localization within the PC. No obvious defects in the structure or distribution of the cilia were observed in the transgenic embryos, and IFT88, a key component responsible for intraflagellar transport, was normally expressed in the cilia of *sx1002* embryos (Fig. S1B), indicating that Nphp3N-AKAR2-CR expression does not affect the structural and functional integrity of PC.

To determine whether changes in PKA activity can be detected in the *sx1002* embryos, embryos were treated with FSK and the relative FRET efficiency assayed. Consistent with the results in NIH3T3 cells, FSK treatment resulted in a marked increase in the FRET efficiency in *sx1002* embryonic PC (Fig. 1I and 1J). These results indicate that the new ciliary PKA biosensor we have developed can report increase in PKA activity in the PC of cells and whole organisms. To test the sensitivity of the Nphp3N-AKAR2-CR biosensor further, we evaluated the ability of the *sx1002* PKA biosensor in response to a gradient of FSK concentrations. As shown in Figure 1H, FRET efficiency increased progressively with increasing FSK concentrations, with a subtle yet discernible change observed even at 1 µM FSK, indicating that the ciliary PKA biosensor exhibits high sensitivity in detecting minute alterations of PKA within the PC. Notably, although all tested FSK concentrations increased FRET efficiency in the PC, only 200 µM FSK treatment, resulting in a ∼7% increase in FRET efficiency, fully phenocopied the effects of HH pathway inhibition by CyA, as revealed by the absence of both Prox1a+ and En2a+ muscle cells in treated embryos (Fig. S2). By contrast, 1 µM and 10 µM FSK elicited an increase, albeit modest (1∼3%) in FRET efficiency within the PC without significantly altering HH activity.

Next, we asked whether genetic manipulation of PKA activity can be detected by the ciliary PKA biosensor. Injection of *dnPKA* mRNA into newly fertilized eggs led to a significant decrease in the FRET efficiency in the PC of resulting embryos compared to controls (Fig. 1I and 1J), Unexpectedly, however, and contrary to the effects of FSK described above, no significant change of FRET was observed in embryos injected with *cPKA* mRNA (Fig. 1I and 1J). We surmised that this could be due to the inability of the injected cPKA to localize to the PC. To test this hypothesis, the subcellular localization of cPKA and dnPKA was visualized by expressing fluorescently tagged versions of each protein (cPKA-eGFP and dnPKA-eGFP) in zebrafish embryos. IF staining of 18hpf embryos revealed that cPKA-eGFP was mainly localized in the cytoplasm, whereas the dnPKA-eGFP was observed both in the cytoplasm and the PC (Fig. 2B and 2D). These results indicate that our biosensor is sensitive to PKA activity specifically in the PC.

**Figure 2.**
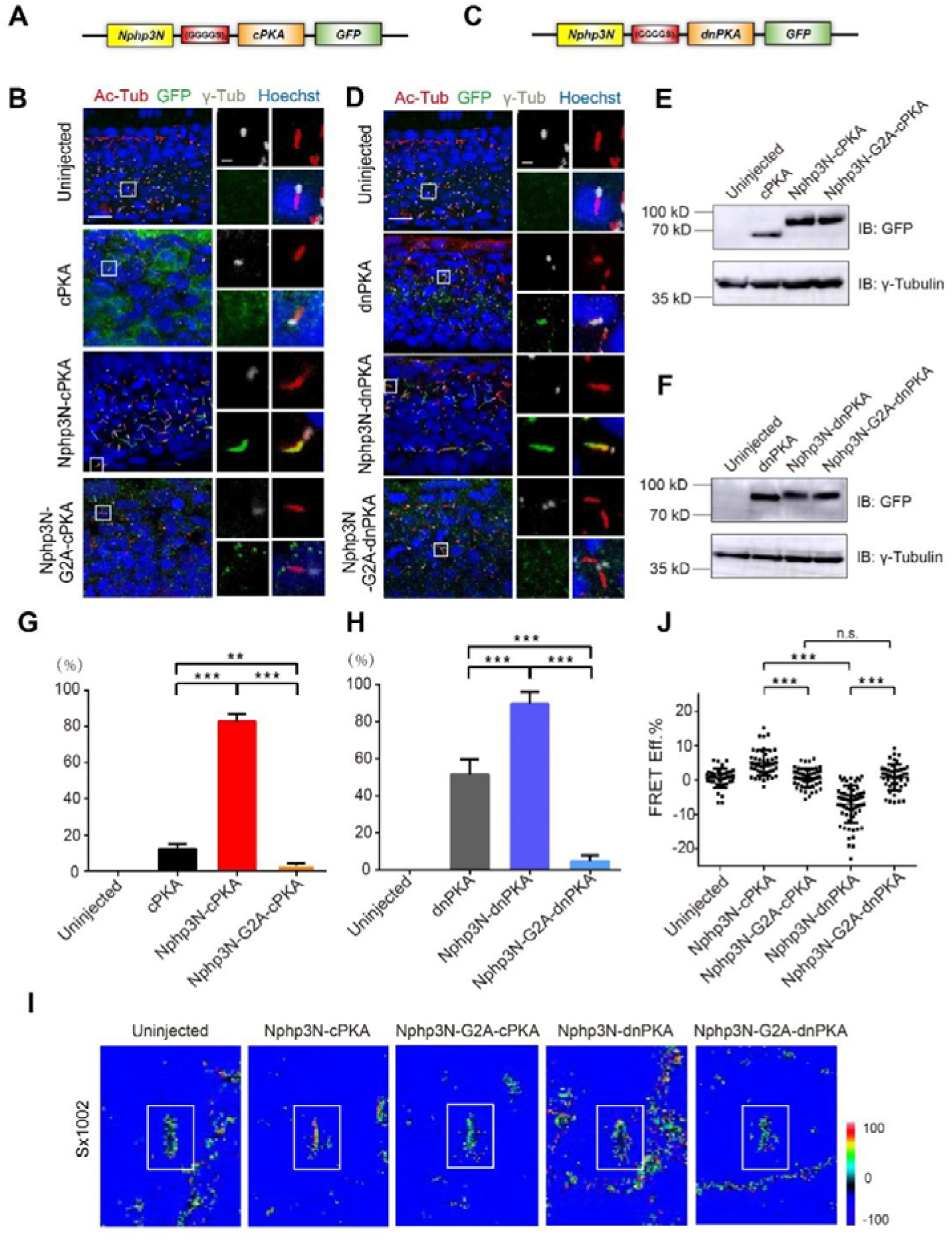
Nphp3N efficiently targets cPKA and dnPKA to the PC. (A and C) Schematic diagram of the linker-modified versions of the *Nphp3N-cPKA/dnPKA-eGFP* vector. (B and D) Subcellular localization of the cPKA/dnPKA-eGFP, Nphp3N(WT/G2A)-cPKA/dnPKA-eGFP in embryos at 18 hpf, as indicated by eGFP in green. The cilia were labelled by Ac-Tub in red, the basal body by γ-tubulin in gray, and the nuclei by Hoechst in blue. Frames on the right panel indicates cilia depicted in the inset, with the basal body on upper left, cilia on upper right, indicated cPKAs/dnPKAs on lower left, and merge images on lower right. Scale bars, 10 μm for each left panel and 2.5 μm for the right panel. (E and F) Immunoblot of lysates from 18hpf zebrafish embryo expressing indicated GFP-tagged forms of cPKA/dnPKA and Nphp3N(WT/G2A)-cPKAs/dnPKAs. The γ-tubulin was used as loading control. (G and H) Quantification of ciliary colocalization from experiments presented in B and D (*n* = 60 cilia in 3 embryos). One-way ANOVA was used for analysis, and *** indicates ‘significant’ with *P*<0.001, ** indicates ‘significant’ with *P*<0.01. (I) The FRET ratio image of ciliary AKAR2-CR in *sx1002* expressing Nphp3N(WT/G2A)-cPKAs/dnPKAs. Representative images of the FRET ratio for each condition in the pseudocolor scale. (J) Statistical analysis of the ciliary FRET efficiency in *sx1002* expressing Nphp3N(WT/G2A)-cPKAs/dnPKAs ^(n^uninjected^=37 cilia, n^Nphp3N^-^cPKA^=46 cilia, n^Nphp3N-G2A-cPKA^=49 cilia, n^Nphp3N^-dnPKA^=54 cilia, n_Nphp3N-G2A-cPKA_=46 cilia). One-way ANOVA was performed, and the ns indicates ‘not significance’ with P>0.05, while *** indicates ‘significant’ with *P*<0.001.

### Specific delivery of functional PKA to the primary cilia

Given the observed divergence of the subcellular localization of the exogenous subunits, we set out to test the cilia specificity of PKA activity further, by targeting both cPKA and dnPKA to the PC using Nphp3N CTP. To avoid structural constraints and misfolding of the two fused functional polypeptides, a hydrophilic flexible linker peptide (GGGGS)_2_ was inserted, thereby increasing the distance between Nphp3N CTP and the PKA functional domain and thus enhancing the proper folding of each domain (Fig. 2A and 2C). The Nphp3N-cPKA and dnPKA fusions were expressed normally in 18hpf embryos (Fig. 2E and 2F) and both localized specifically in PC, with the localisation efficiencies up to 82.5% and 89.5%, respectively (Fig. 2B, 2D, 2G and 2H). Nphp3N is also capable of targeting cPKA/dnPKA to cilia in NIH3T3 cells (Fig. S3). The Glycine of the second residue of Nphp3N is critical for its ciliary targeting function, and the G2A mutation impairs its ciliary localization property (Nakata et al., 2012; Zhang et al., 2022). Consistently, Nphp3N-G2A-cPKA and dnPKA fusion proteins were observed to localize only in the cytoplasm and not in the shaft of the PC (Fig. 2B, 2D, 2G and 2H), thus providing an effective means of comparing cilia-versus cyto-targeted PKA function.

To investigate the activity of ciliary localized Nphp3N-cPKA/dnPKA, mRNAs for either construct were injected into embryos of the *sx1002* PKA biosensor line. As expected, the cilia-cPKA and dnPKA showed an increase or decrease, respectively, in the FRET efficiency of ciliary AKAR2 (Fig. 2I and 2J). By contrast, no change in FRET was detected following injection of the cyto-cPKA/dnPKA mRNAs compared to the control group (Fig. 2I and 2J). Taken together, these results indicate that the newly established transgenic ciliary PKA biosensor line *sx1002* can effectively report changes in PKA activity within PC.

### PKA acts in the PC to regulate HH pathway activity

Previously, Truong *et al*. employed RAB23 variants to target dnPKA to the interior and exterior of cilia and concluded that only ciliary dnPKA could specifically upregulate the Hh pathway in zebrafish embryos (Truong et al., 2021). However, the activation of HH pathway achieved by this targeted dnPKA was much weaker than that resulting from expression of untargeted dnPKA, as assayed by the expansion of Prox1a and En2a expression in skeletal muscle cells. To investigate this disparity further, we introduced our Nphp3N(G2A), dnPKA constructs into embryos of a newly established transgenic line, *sx1005* (see Materials and Methods), carrying a previously described *En2a::GFP* reporter construct (Maurya et al., 2013). In contrast to RAB23-Q68L-targeted cilia-dnPKA, Nphp3N-dnPKA significantly increased the number of En2a/Prox1a double-positive muscle cells, consistent with the previously described hyper activation of HH pathway effected by injecting dnPKA (Hammerschmidt et al., 1996; Wolff et al., 2003), whereas Nphp3N(G2A)-dnPKA had virtually no effect on HH pathway activity (Fig. 3B and 3C). Consistently, only Nphp3N-dnPKA induced ectopic upregulation of the Hh pathway target genes, such as *ptch2*, *oligo2*, and *nkx2.2* as detected by *in situ* hybridisation, indicating that cilia-dnPKA mediated by Nphp3N specifically activates the HH pathway, significantly more effectively than the cilia-dnPKA delivered via RAB23-Q68L (Fig. 3D).

**Figure 3.**
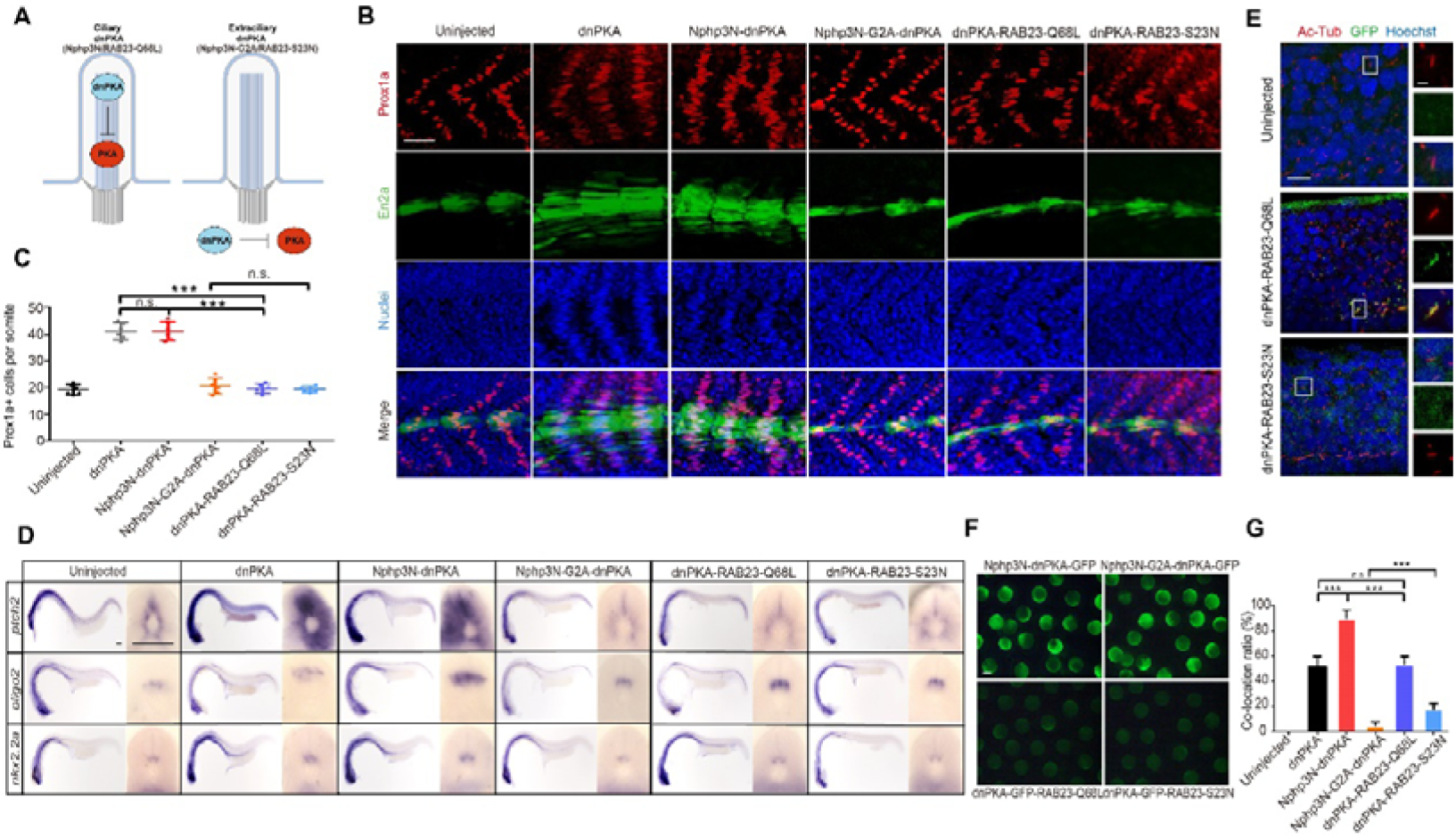
Nphp3N drove cilia localization of the dnPKA regulates the HH pathway more effectively. (A)Schematic digram of dnPKA at distinct subcellular locations in this assay. (B) The activity of Hh pathway was indicated by expression of Prox1a in red and En2a:eGFP in green in the embryos of Tg (*en2a*:*eGFP*)^sx1005^ expressing indicated Nphp3N (WT/G2A)-dnPKA and RAB23 (Q68L/S23N)-dnPKA, respectively. The nuclei were labelled by Hoechst in blue. Scale bar, 50 μm. (C) Quantification of Prox1a+ cells from experiments presented in B (*n* = 6 somites from 3 embryos). One-way ANOVA was performed. And ns indicates ‘not significant’ with *P*>0.05, *** indicates significant with *P*<0.001. (D) *In situ* hybridization of *ptch2*, *nkx2.2a* and *olig2* on the 24 hpf embryos expressing indicated dnPKAs. Each panel showed a full view of the embryo on the left and a cross-sectional view of a somite on the right (*n* = 3 for each sample). Scale bars, 100 μm. (E) Subcellular localization of the dnPKA-eGFP-Rab23 Q68L/S23N in embryos at 18 hpf, as indicated by eGFP in green. The cilia were labelled by Ac-Tub in red, and the nuclei by Hoechst in blue. Frames on the right panel indicates cilia depicted in the inset, with cilia on the top, indicated cPKAs/dnPKAs in the middle, and merge images at the bottom. Scale bars, 10 μm for the left panel and 2.5 μm for the right panel. (F) Transient expression of the indicated Nphp3N (WT/G2A)-dnPKA and RAB23 (Q68L/S23N)-dnPKA in zebrafish embryos at 6 hpf were indicated by eGFP in green. Scale bars, 500 μm. (G) Quantification of ciliary colocalization from experiments presented in Fig 2D and E (*n* = 60 cilia in 3 embryos). One-way ANOVA was used for analysis, and *** indicates ‘significant’ with *P*<0.001, ns indicates ‘not significant’ with *P*>0.05.

We surmised that this reflects the low expression and/or ciliary targeting efficiency of RAB23-Q68L. Accordingly, we compared the efficiency of the two ciliary delivery systems: the RAB23-Q68L-dnPKA and RAB23-S23N-dnPKA constructs were generated and their expression and subcellular localization examined by mRNA microinjection into zebrafish embryos. The dnPKA-eGFP fused with RAB23 variants was expressed at much lower levels than the same protein fused with Nphp3N and its variants at 6 hours post injection (hpi) and later stages (Fig. 3F). Similarly, the efficiency of ciliary localization of the dnPKA-eGFP fused to RAB23-Q68L was much lower than of the same protein fused to Nphp3N (Fig. 3E and 3G).

Although previous studies have confirmed that the RI subunit of PKA is localized to PC (Mick et al., 2015), the ciliary localization of PKA-C under normal physiological conditions remains unclear (Truong et al., 2021; Walker et al., 2024). To investigate whether ciliary localization of PKA-C is sufficient for modulation of HH pathway activity, we used the Nphp3N signal peptide to express cPKA specifically in the PC of the *sx1005* transgenic embryos. Injection of mRNA encoding such cilia-targeted cPKA resulted in a significant decrease of En2a+ and Prox1a+ muscle cells in *sx1005* embryos (Fig. 4B and 4C). Consistently, the expression of *ptch2*, *oligo2*, and *nkx2.2* was reduced in these embryos (Fig. 4D). By contrast, embryos injected with either untargeted cPKA or the cyto-cPKA, showed normal expression of HH-pathway target genes and similar numbers of the En2a+ and Prox1a+ muscle cells to those in wild-type embryos (Fig. 4B, 4C and 4D). Taken with the finding that the untagged cPKA localizes mainly in the cytoplasm (Fig. 2B), this indicates that the HH signaling pathway is inhibited only by PKA activity in the PC. We also generated a RAB23-tagged form of cPKA. Consistent with our findings with the RAB23-dnPKA, RAB23-cPKA exerted no significant effect on HH signaling, most likely due to its low expression and sub-optimal ciliary targeting efficiency (Fig. S4).

**Figure 4.**
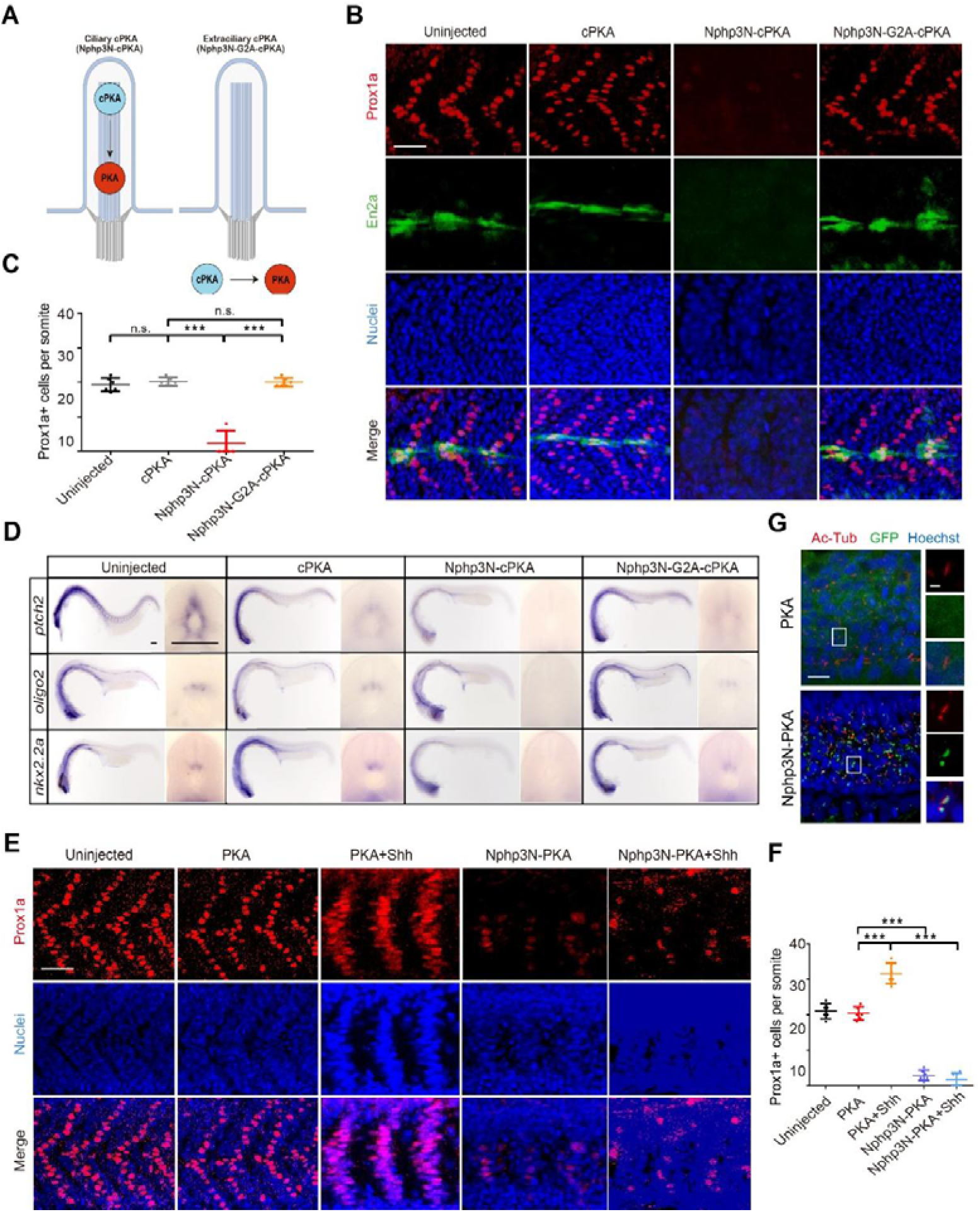
Only cilia-targeted cPKA can modulate HH activity. (A)Schematic digram of cPKA at distinct subcellular locations in this assay. (B) The activity of Hh pathway was indicated by expression of Prox1a in red and En2a:eGFP in green in the embryos of Tg (*en2a*:*eGFP*)^sx1005^ expressing indicated Nphp3N (WT/G2A)-cPKA. The nuclei were labelled by Hoechst in blue. Scale bar, 50 μm. (C) Quantification of Prox1a+ cells from experiments presented in B (*n* = 6 somites from 3 embryos). One-way ANOVA was performed. And ns indicates ‘not significant’ with *P*>0.05, *** indicates significant with *P*<0.001.(D) *In situ* hybridization of *ptch2*, *nkx2.2a* and *olig2* on the 24 hpf embryos expressing indicated cPKAs. Each panel showed a full view of the embryo on the left and a cross-sectional view of a somite on the right (*n* = 3 for each sample). Scale bars, 100 μm. (E) The activity of Hh pathway was indicated by expression of Prox1a in red in the embryos of Tg (*en2a*:*eGFP*)^sx1005^ expressing PKA, PKA+SHH, Nphp3N-PKA or Nphp3N-PKA+SHH. The nuclei were labelled by Hoechst in blue. Scale bar, 50 μm. (F) Quantification of Prox1a+ cells from experiments presented in E (*n* = 6 somites in 3 embryos). One-way ANOVA was used for analysis, ns indicates ‘not significant’ with *P*>0.05, *** indicates ‘significant’ with *P*<0.001. (G) Subcellular localization of the PKA and Nphp3N-PKA in embryos at 18 hpf, as indicated by eGFP in green. The cilia were labelled by Ac-Tub in red, and the nuclei by Hoechst in blue. Frames on the right panel indicates cilia depicted in the inset, with the cilia on the top, the indicated PKA in the middle, and merge layer at the bottom. Scale bars, 10 μm and 2.5 μm (inset).

The failure to detect PKA-C in the PC by immunofluorescence in the physiological state suggests that the low level of PKA-C must be strictly controlled (Fig. 4G). If so, increasing the levels of PKA-C in the PC should have a similar effect as the cilia-cPKA. To test this hypothesis, Nphp3N-PKA-C mRNA was injected into embryos. Like the cilia-cPKA, the protein localized to the PC (Fig. 4G) and efficiently suppressed HH activity, as shown by the depletion of both Prox1a+ and En2a+ muscle cells in the myotome (Fig. 4E and 4F). In contrast, the untagged PKA-C, which was only observed in the cytoplasm, had no effect on HH activity (Fig. 4E and 4F).

### Cilia targeting of PKA is essential even in the absence of PC

In vertebrates, the dependence of HH signaling upon the PC is reflected by disruption of HH pathway activity caused by PC aberrations in mouse and zebrafish embryos (Huangfu et al., 2003; Bazzi et al., 2014; Davey et al., 2006). Our findings imply that HH pathway activity is efficiently suppressed by a basal level of PKA-C activity within the PC. Accordingly, loss of PC integrity might lead to suppression of pathway activity being mediated, albeit less efficiently, by cytoplasmic PKA. To investigate the regulation of the HH pathway by ciliary PKA under conditions of ciliary dysfunction, the PC was disrupted in zebrafish embryos by depleting both maternal and zygotic Kif3a, one of critical factor for primary ciliogenesis (Corbit et al., 2008; Li et al., 2020; Cullen et al., 2021; Xie et al., 2022). As previously shown (Xie et al., 2022), *MZkif3a* embryos show a reduction of the number of PCs in neural tube and otic vesicle cells (Fig. S5). Since this mutant line is a selected transgenic strain by both red and green fluorescence [the offspring of Tg(*kop::cas9-p2a-egfp-UTRnanos3*) and Tg(*3×sgRNA-kif3a-mcherry*)], to avoid interference from overlapping fluorescent signals, the following staining of Prox1a- and En2a-positive cells had to be separately detected.

Compared with wild-type embryos, the number of Prox1a+ muscle cells in *MZkif3a* mutants increased slightly, while the number of En2a+ muscle cells was significantly increased, which is typical of mild upregulation of the HH pathway in cilia defective mutants (Fig. 5A and 5C). Injection of cyto-cPKA showed little effect in *MZkif3a* mutants (Fig. 5A and 5C). By contrast, injection of cilia-cPKA into *MZkif3a* mutants resulted in a dramatic reduction in the number of En2a+ muscle cells, similar to that observed in wild-type embryos injected with cilia-cPKA (Fig. 5D). Consistently, the number of Prox1a+ cells was also greatly reduced (Fig. 5B). These findings indicate that cilia-cPKA inhibits HH pathway activity even in the absence of PC, albeit with a slightly attenuated effect compared to identically treated wild-type embryos. Conversely, injection of cilia-dnPKA into *MZkif3a* mutants led to an increase in Prox1a+ and En2a+ muscle cells (Fig. 5B and 5D), comparable to that observed in wild-type embryos, whereas cyto-dnPKA had little effect (Fig. 5A and 5C). These results indicate that only ciliary PKA, but not cytoplasmic PKA, can significantly regulate HH pathway activity even in the absence of PC. Notably, neither the ciliary cPKA nor the ciliary dnPKA can fully deplete or increase the numbers of Prox1a+ cells in the absence of PC as they do in wild-type embryos, indicating that the regulatory role of PKA in the HH pathway is optimally PC-dependent. Notably, by subcellular localization analysis, we observed that in *MZkif3a* mutant, cilia-cPKA/dnPKA tended to aggregate in puncta close to the basal body, while the cyto-cPKA/dnPKA displayed ubiquitous expression (Fig. 5E). This suggests that, in the absence of cilia, ciliary-targeted cPKA and dnPKA may function within a cilia-related compartment near the basal body and that such sub-cellular localization is critical for PKA-mediated regulation of the HH pathway.

**Figure 5.**
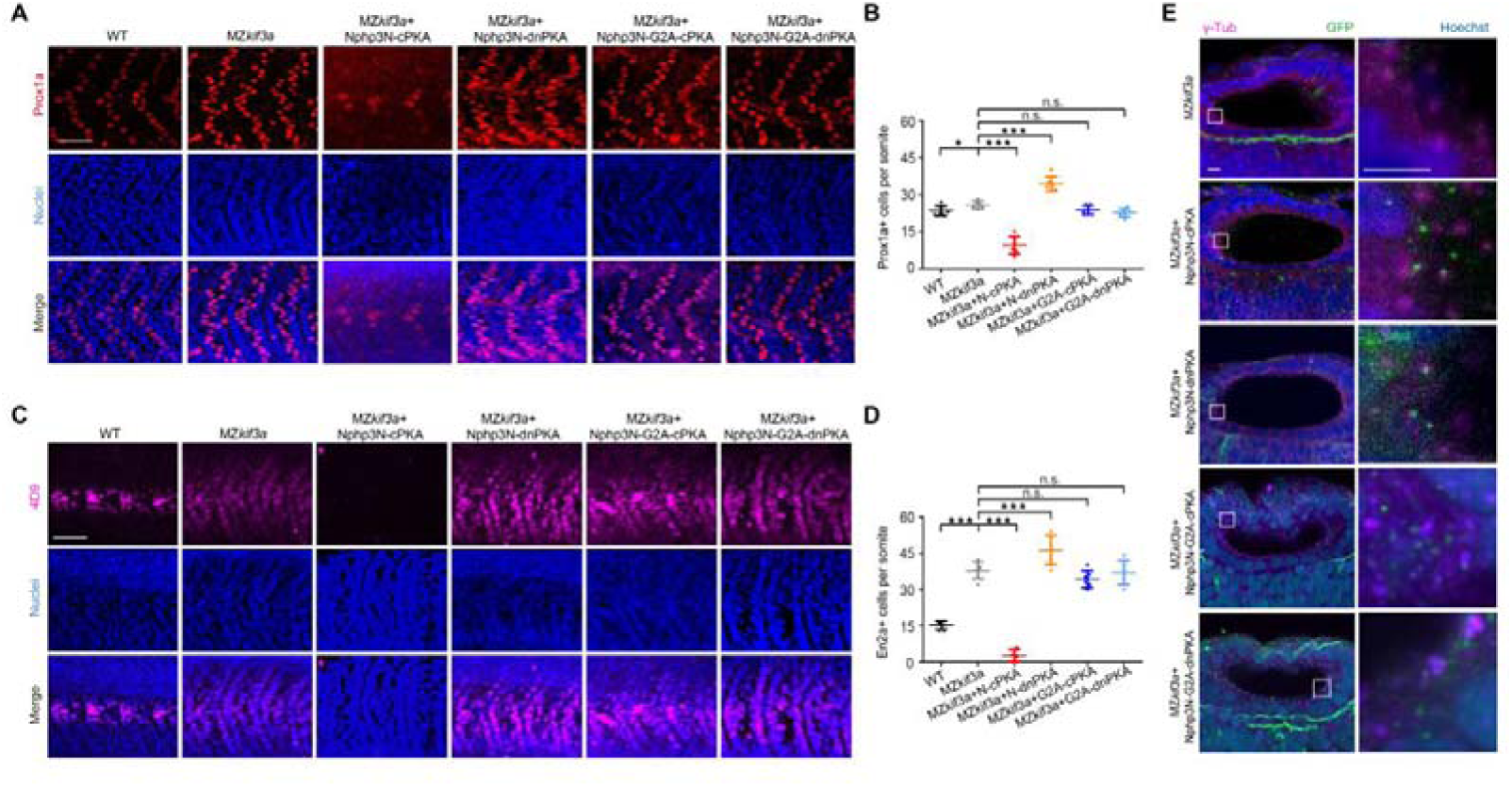
Localisation of PKA to the basal body is key to its regulation of HH. (A) Immunofluorescence staining images of Prox1a (red) in MZ*kif3a* mutants expressing Nphp3N (WT/G2A)-cPKA/dnPKA. (B) Quantification of Prox1a+ cells from experiments presented in A (*n* = 6 somites from 3 embryos). One-way ANOVA was performed. And ns indicates ‘not significant’ with *P*>0.05, * indicates significant with *P<*0.05, *** indicates significant with *P*<0.001. (C) Immunofluorescence staining images of En2a (magenta) in MZ*kif3a* mutants expressing Nphp3N (WT/G2A)-cPKA/dnPKA. (D) Quantification of En2a+ cells from experiments presented in C (*n* = 6 somites from 3 embryos). One-way ANOVA was performed. And ns indicates ‘not significant’ with *P*>0.05, *** indicates significant with *P*<0.001. (E) Subcellular localization of the Nphp3N(WT/G2A)-cPKA/dnPKA in MZ*kif3a* at 24 hpf, as indicated by eGFP in green. The basal bodies were labelled by gamma tubulin in magenta, and the nuclei by Hoechst in blue. The right panel shows the basal bodies and PKA variants in the framework. Scale bars, 10 μm and 2.5 μm (inset).

### Overexpression of SMO fails to fully activate the HH pathway through complete inhibition of PKA activity

Having established that HH pathway activity is modulated specifically by PKA localised to the PC, we next investigated the effect of HH pathway activation on PKA. First, NIH3T3 cells transiently expressing Nphp3N-AKAR2-CR were treated with the SMO agonist SAG or the antagonist, cyclopamine (CyA). Unexpectedly, although both drugs significantly altered HH pathway activity as indicated by changes in expression of the HH pathway target genes *gli1* and *ptch1* in transfected cells (Fig. S6A), we failed to detect any change of FRET efficiency within the cilia following either treatment. (Fig. 6A and 6B). Next, we assayed the ciliary PKA activity in *sx1002* zebrafish embryos injected with *shh* mRNA or treated with CyA. As previously reported (Wolff et al., 2003), these treatments resulted in an increase or decrease, respectively, in the number of Prox1a+ muscle cells (Fig.S6B and S6C). However, as with the SAG and CyA treated NIH3T3 cells, no significant change in PKA activity in the PC was detected in either case (Fig. 6C and 6D).

**Figure 6.**
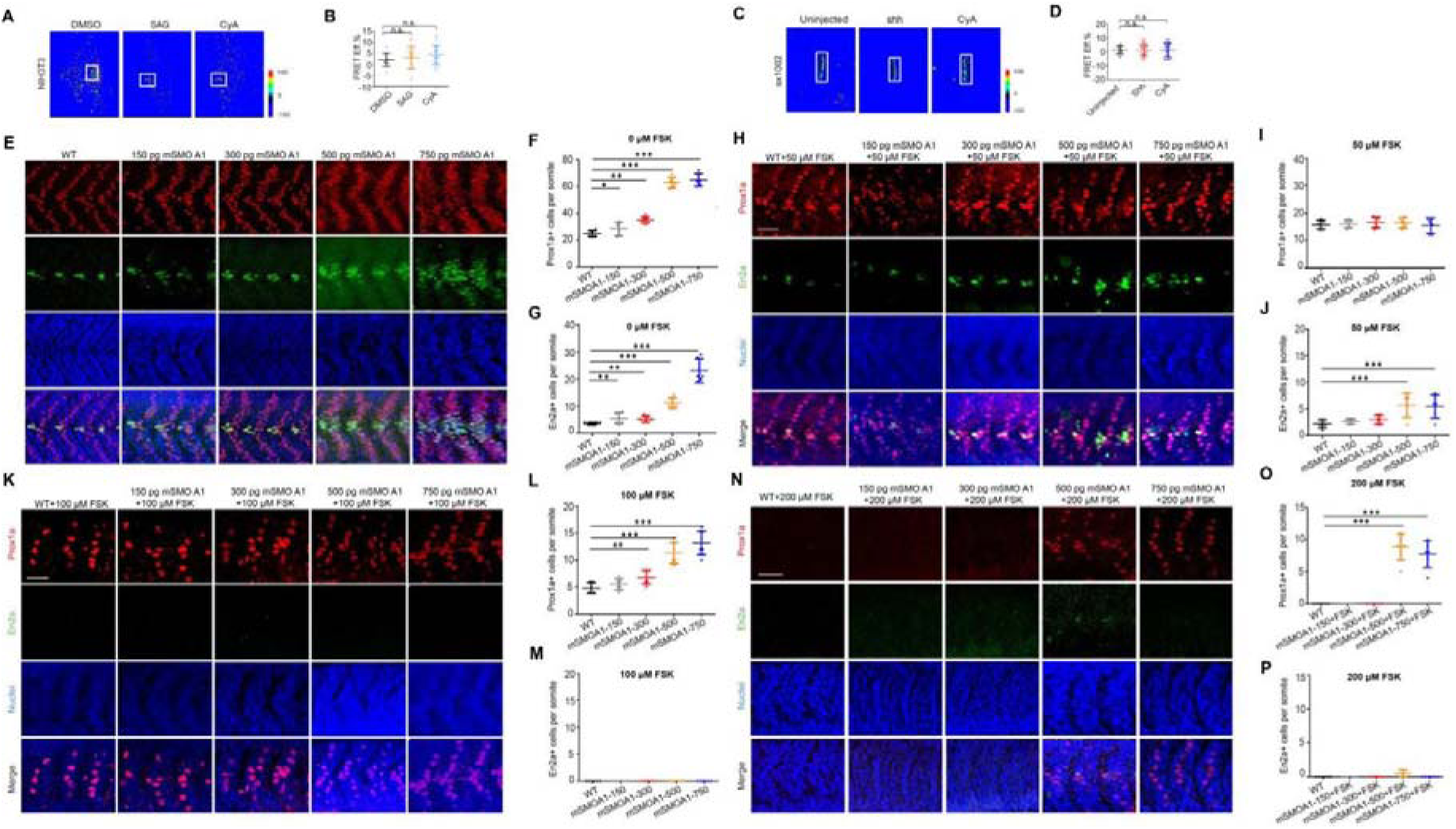
SMO may not control HH pathway activity exclusively by modulation of PKA (A) The FRET ratio image of ciliary AKAR2-CR in NIH3T3 cells treated with DMSO, 1 μM SAG and 5 μM cyclopamine (CyA), respectively. (B) Statistical analysis of the ciliary FRET efficiency in NIH3T3 cells in response to DMSO, SAG and CyA, respectively (n_DMSO_=12 cilia, n_SAG_=20 cilia n_CyA_=22 cilia). One-way ANOVA was performed, and the ns indicates ‘not significant’ with *P*>0.05. (C) The FRET ratio image of ciliary AKAR2-CR in *sx1002* injected with *shh* mRNA or treated with CyA. (D) Statistical analysis of the ciliary FRET efficiency in *sx1002* treated with *shh* and CyA (n_uninjected_=18 cilia, n_shh_=33 cilia, n_cyA_=31 cilia). One-way ANOVA was performed, and the ns indicates ‘not significant’ with *P*>0.05.(E, H, K, N) Immunofluorescence staining images of Prox1a (red) and En2a (green) in wild-type and those injected with a concentration gradient of mSMOA1 under treatment with varying concentrations of FSK. E, 0 μM FSK. H, 50 μM FSK. K, 100 μM FSK. N, 200 μM FSK. The nuclei were labelled by Hoechst in blue. Scale bar, 50 μm. (F, I, L, O) Quantification of Prox1a+ cells from experiments presented in E, H, K, N (*n* = 6 somites from 3 embryos). One-way ANOVA was performed. ‘**’ indicates ‘significant’ with *P*<0.01. ‘***’ indicates ‘significant’ with *P*<0.001. (G, J, M, P) Quantification of En2a+ cells from experiments presented in E, H, K, N (*n* = 6 somites from 3 embryos). One-way ANOVA was performed. ‘**’ indicates ‘significant’ with *P*<0.01. ‘***’ indicates ‘significant’ with *P*<0.001.

These findings stand in contrast to the proposal that activated SMO drives the response to HH signals by directly sequestering PKA-C within PC via its PKI-like domain (Walker et al., 2024; Nguyen et al., 2025; Steiner et al., 2025). If this were the case, activated SMO and the ciliary PKA should show a stoichiometric binding relationship upon HH pathway activation. To test this hypothesis, we performed a dose-response titration assay of the constitutively active SMO variant (SMOA1) (Taipale et al., 2000) against FSK in zebrafish embryos. First, we analyzed muscle cell differentiation phenotypes in zebrafish embryos treated with a range of FSK concentrations and SMOA1. As shown in Supplementary Figure S2, 10 µM FSK induced a very slight reduction in Prox1a+ muscle cell number, whereas 100 µM FSK significantly inhibited the HH pathway, as manifest by a loss of En2a+ and a marked decrease in Prox1a+muscle cells. At 200 µM, FSK exerted a CyA-like effect, completely eliminating HH-dependent cell types. This result indicates that at this concentration range, FSK (and by extension, PKA) and the HH pathway activity display a negative linear relationship (Fig. S2). Thus, we selected a 50∼200µM range of FSK for subsequent titration experiments. Parallel concentration-testing of SMOA1 revealed that at doses below 500pg, the number of Prox1a+ and En2a+ cells increased in a dose-dependent manner (Fig. 6E). However, SMOA1 expression within this range is insufficient to activate the HH pathway fully, while, at doses of 500pg or higher, a robust increase in Prox1a+ and En2a+ muscle cells was observed, the number of double-positive cells peaking at 750pg (Fig. 6F and 6G), indicating excessive activation of the HH pathway under these conditions. Embryos injected with 1000pg of SMOA1 showed severe deformities, most likely reflecting the toxicity of high concentration of mRNA (not shown). Accordingly, we used a range of 150-750pg for the titration experiments.

Treatment with 200 µM FSK resulted in a complete suppression of double-positive cells in all SMOA1 dose groups compared to the untreated control group (Fig. 6N). When the dose of SMOA1 exceeded 500pg, a few Prox1a+ muscle cells were present (no more than 10 per somite) (Fig. 6O), whereas En2a+ muscle cells were completely absent (Fig. 6P). ThusSMOA1 overexpression failed to reverse the FSK-mediated inhibition of the HH pathway. To rule out the possibility that non-physiological levels of PKA activity caused by excessively high FSK concentrations prevented SMOA1 from rescuing FSK-induced HH pathway inhibition, we reduced the concentration of FSK, while varying the expression dose of SMOA1. Under 100 µM FSK treatment, the HH pathway was sub-maximally inhibited (Fig. 6K): En2a+ muscle cells were undetectable (Fig. 6M), while the number of Prox1a+ muscle cells decreased sharply. A slight recovery in the number of Prox1a+ muscle cells was observed in embryos injected with 500pg and 750pg SMOA1 (Fig. 6L) but no rescue of En2a+ cellswas evident (Fig. 6M). At 50µM FSK only a small increase in En2a+ cell number was seen in the 500pg and 750pg SMOA1 injection groups (Fig. 6H and 6J), while there was no significant difference in the number of Prox1a+ muscle cells among all groups (Fig. 6H and 6I). These results indicate that PKA-mediated inhibition of the HH pathway cannot be fully relieved by SMO activation and perturbations in SMO levels can only modulate PKA activity within a low-threshold range.

## Discussion

Primary cilia (PC) play important roles in regulating vertebrate HH signal transduction and PKA activity in the PC is considered to be a critical aspect of the requirement for the PC in this signaling cascade; however, the precise details of the processes supported by the PC are still unclear. Compartmentalized activity underpins the precise regulation of key cellular processes mediated by PKA (Li et al., 2013; Wong et al., 2004; Taylor et al., 2012). Tracking the activity of PKA in living cells enhances our understanding of this regulation. To achieve this, two strategies could be adopted: measuring either the cAMP concentration or PKA activity directly. Tools for monitoring cAMP level in PC by Arl13b-H187 or 5HT6-mCherry-cADDis have been reported (Tewson et al., 2016; Moore et al., 2016; Jiang et al., 2019). Although the major pathway activated by cAMP is PKA-dependent, cAMP can also activate Epac and cyclic nucleotide-gated (CGN) ion channels directly, independently of PKA (Beavo and Brunton, 2002). Thus, cAMP level is not synonymous with PKA activity.

To monitor PKA activity in the PC, Moore *et al*. previously utilized the ciliary GPCR 5HT_6_, to target another FRET-based PKA sensor, AKAR4, which reported slightly higher PKA activity in cilia compared to cytoplasm (Moore et al., 2016). However, this approach suffers from the possibility that as a typical ciliary GPCR, the 5HT_6_, could itself modulate PKA activity by coupling to G_αs_, contributing to the increase in ciliary PKA activity.

To circumvent these limitations, we have developed an improved sensor for monitoring PKA activity in the PC, both in tissue culture and in living embryos. The Nphp3N CTP targeting peptide that we used is much smaller than Arl13b or 5HT_6_ and has minimal effect on PC integrity (Zhang et al., 2022). In addition, the AKAR2-CR, which uses the Clover-mRuby as the FRET pair shows higher sensitivity and stability than the AKAR4 that uses the Cerulean-cpVenus as the FRET pair (Demeautis et al, 2017). We generated a stable zebrafish transgenic line ubiquitously expressing the Nphp3N-AKAR2-CR fusion protein, allowing monitoring of PKA activity at the individual PC level. A key advantage of this biosensor is its utility for detecting subtle fluctuations in ciliary PKA activity, which outperforms previous tools such as CREB-based biosensors, limited to global cellular readouts (Arveseth et al., 2021). The novel biosensor detected a basal PKA activity, as evidenced by reduced FRET efficiency upon dnPKA expression, and dose-dependent increases with FSK treatment. The level must be strictly controlled as we also failed to detect cPKA within the PC when it was overexpressed. Using our biosensor, we found that PKA activity in the PC remained largely unchanged following perturbation of the HH pathway at the level of SMO. Furthermore, we demonstrate that only ciliary-targeted PKA, and not its cytoplasmic counterpart, modulates HH signaling, even in the absence of intact cilia, where targeted PKA aggregates near the basal body. These findings underscore the compartmentalized nature of PKA function in HH repression and suggest that a low basal level of ciliary PKA maintains the pathway in an OFF state.

Recently, the use of an alternative ciliary PKA biosensor, based on detection of the pVASP by immunofluorescence staining, in both zebrafish embryos and NIH3T3 cells has been reported (Nguyen et al., 2025). This PKA biosensor displays very highly sensitivity in cilia responding to concentrations of FSK as low as 100nM. However, unlike our FRET-based Nphp3N-AKAR2-CR sensor, the Arl13b-GFP-VASP sensor does not allow monitoring of PKA activity in real-time. Moreover, increasing the levels of Arl13b in the PC could cause SMO and GLI2 accumulation independent of exogenous SHH ligand (Shi et al., 2023; Gigante et al., 2020). These factors need to be borne in mind when considering the differences in findings using the two sensors.

Although the role of PKA in HH signal transduction is well established, it is still controversial whether PKA regulation of the vertebrate HH pathway is absolutely dependent on the PC. While a weak signal of the PKA-C has been detected in PC 6 hours after HH pathway activation with the SMO agonist SAG (Walker et al., 2024), the ciliary localization of PKA under basal conditions has not been reported. Moreover, rather than localization to the PC, PKA has been reported to localize specifically at the basal body, which suggested that this basal body localization of PKA is critical for its role in the HH pathway (Tuson et al., 2011; Peng et al., 2021). To resolve the subcellular site of PKA action, Troung et al. used RAB23 variants to deliver dnPKA to PC and demonstrated that cilia-dnPKA could specifically activate the HH pathway (Troung et al., 2021). However, the magnitude of HH activation by ciliary dnPKA in these experiments was modest. Nevertheless, this spatially restricted manipulation strategy provided a proof of concept for dissecting PKA compartment-specific function. Building on this foundation, ciliary-targeted cPKA and dnPKA using Nphp3N with a (GGGGS)_2_ linker, achieved a high PC-localisation efficiency (82.5∼89.5%) compared to the lower expression and targeting efficiency of the RAB23 variants. This superior Nphp3N/G2A mediated targeting enabled precise manipulation. Only ciliary cPKA/dnPKA altered FRET in *sx1002* and HH outputs in *sx1005* embryos, depleting or expanding En2a+/Prox1a+ muscle cells and targets like *ptch2*, *oligo2*, *nkx2.2*. Cytoplasmic variants had no effect, underscoring spatial specificity of PKA in the regulation of HH pathway. This aligns with compartmentalized PKA in vertebrates, where ciliary PKA phosphorylates Gli2/3 at the base, promoting repressor forms (Li et al., 2017; Zhou and Jiang, 2022). Strikingly, in MZ*kif3a* mutants lacking most PCs, ciliary-targeted PKA still modulated HH activity albeit attenuated, and aggregated near basal bodies. This mild HH upregulation in mutants was suppressed or enhanced only by ciliary targeted cPKA or dnPKA, while cytoplasmic forms had no effect. Basal body localization suggests that a periciliary compartment, possibly the ciliary pocket or transition zone, serves as a signaling nexus when cilia are absent (May-Simera and Kelley, 2012). This extends the established paradigm of ciliary dependence in vertebrate HH signaling, where defects mildly activate the pathway, to reveal the adaptive capacity of PKA in sustaining HH pathway regulation under ciliary perturbation. Importantly, despite the absence of detectable PKA-C localization in PC, perturbation of ciliary PKA activity at a certain threshold, as shown by Nphp3N-cPKA and dnPKA (but not RAB23-cPKA/dnPKA) dramatically changed the HH pathway activity in our assay. This indicates that the maintenance of the basal level of ciliary activity is critical for the ON and OFF states in vertebrate HH pathway. This paradox, undetectable PKA-C but measurable activity, suggests tight regulation, possibly via A-kinase anchoring proteins (AKAPs) that tether PKA to the cilia, ensuring localized phosphorylation without bulk accumulation (Kultgen et al., 2002; Bachmann et al., 2016).

While our study confirms that basal ciliary PKA activity is critical for regulating the HH pathway, direct monitoring of PKA activity across distinct HH pathway states remains lacking. When perturbing the HH pathway at the level of SMO, our optimized PKA biosensor showed no change in FRET efficiency in NIH3T3 cells or zebrafish embryos in response either to SAG/Shh or CyA, despite robust HH target gene modulation. There are at least two possible explanations of these results: first, ciliary PKA activity is not significantly changed upon activation or inhibition at the level of SMO; second, the ciliary PKA biosensor is not sensitive enough to detect a subtle PKA change during HH signal transduction. Interestingly, we note that even with the highly sensitive PKA biosensor Arl13b-GFP-VASP, PKA activity in cells treated with SHH/SAG and 100nM FSK was nearly equivalent to or slightly higher than that in control cells (Nguyen et al., 2025). Following the model that SMO directly sequesters PKA-C in the PC during the ON state of HH pathway (Arveseth et al., 2021; Walker et al., 2024; Steiner et al., 2025) and thus blocks FSK-mediated effect, ciliary PKA activity in cells treated with SHH/SAG plus FSK should match that in cells treated with SHH/SAG alone. These data support the first hypothesis that ciliary PKA activity is not significantly changed at least upon modulation of HH pathway at the level of SMO.

Although it has been shown that SMO directly sequesters the ciliary PKA-C upon activation of HH pathway, whether this sequestration is sufficient to activate the pathway is unclear. To address this, it is necessary to examine the stoichiometric balance of interaction between SMO and PKA-C in the PC. In zebrafish, we observed that 200 µM FSK completely inhibited the HH pathway, while 100 µM FSK exerted strong inhibition, allowing only a small number of Prox1a+ muscle cells to differentiate. In contrast, 50µM FSK caused only a slight reduction in the number of Prox1a+ muscle cells, reflecting a mild inhibitory effect on HH pathway activity. Based on these observations, we propose that 50 µM FSK represents a suitable balanced concentration to overcome SMO-mediated activation. At this FSK concentration, if consistent with the SMO-PKA antagonism model, then overexpression of SMO should decrease PKA activity and thereby activate the HH pathway to a measurable extent. However, our titration experiments revealed that low-dose SMO overexpression was insufficient to rescue the FSK-mediated inhibitory phenotype. Even high-dose SMO overexpression (500-750 pg) only resulted in a modest recovery of pathway activity and failed to fully counteract FSK-mediated modulation of HH output. These results imply that activation of the HH pathway is not solely attributable to SMO-dependent PKA inhibition but involves additional mechanisms, potentially including altered GLI dynamics or ciliary trafficking processes (Kim et al., 2009; Tuson et al., 2011).

One achievement of our work is that we have established a powerful specific delivery system of functional proteins, especially cPKA and dnPKA, to the PC. The smallest cilium target sequence, Nphp3N signal peptide, significantly reduced the side effect on cilia caused by the signal peptide, and the linker sequence ensured the proper function of the proteins targeted to the PC. This delivery system provides a great opportunity to intervene specifically and efficiently in the PC functions to uncover the mechanism of the PC-related cellular functions cellular functions and signal transduction. Notably, our experiments have revealed effects of PC-targeted on HH pathway activity. Aberrant activation of the HH pathway underlies the occurrence and progression of several tumors, especially basal cell carcinoma and medulloblastoma as well as gastro-intestinal and other tumors (Berman et al., 2002; Epstein, 2008; Skoda et al., 2018; Walton et al., 2021), making it a potential target for precision therapeutics. Currently, most such drugs specifically target SMO and show various off-target effects as well as drug resistance. The specific pathway downregulation effected by the Nphp3N-cPKA transgene, suggests the potential for developing novel HH therapeutics.

In conclusion, we have established a delivery method that specifically targets cilia, with high efficiency and without loss of functionality of the targeted proteins. This strategy effectively counteracts any impacts of other tags on the structure and function of primary cilia. Targeting PKA activity specifically in the PC demonstrates the presence of a pool of PKA with activity constrained within a basal range within the PC, which is critical for maintaining HH signaling pathway homoestasit. Our results provide new insights into the mechanism of vertebrate HH signal transduction. whilst the new PC delivery system has potential for precise delivery of multiple proteins and genome-wide tracking of cellular activity, incorporating both temporal and spatial dimensions as well as dynamic information, to enhance the mapping of signal transduction networks.

## Material and Methods

### Cell Culture of HEK293T and NIH3T3 and Transfection

The HEK293T and NIH3T3 cells, procured from ATCC, were maintained in Dulbecco’s Modified Eagle’s Medium (DMEM) supplemented with 10% FBS and 10% penicillin-streptomycin, at 37°C with 5% CO_2_. For ciliogenesis, the cells were initially rinsed with PBS and then subjected to serum starvation by culturing in FBS-free medium for 24 h. For transfection, the Lipofectamine 3000 (Thermo Fisher Scientific, L3000015) or the PEI (MedChemExpress, HY-K2014) were used according to the manufacturer’s protocol. To induce or inhibit the HH signaling, the cells were plated and treated with either 1 μM SAG (MedChemExpress, HY-12848) or 5 μM cyclopamine (MedChemExpress, 4449-51-8), respectively, and then subjected to the following serum starvation for 24 h prior to collection.

### Zebrafish husbandry

Adult zebrafish of the AB strain, sourced from the China Zebrafish Resource Center, were housed at 28 on a 14 h light/10 h dark cycle within the zebrafish facility of Shanxi University. Embryos were obtained from natural mating and grew in E3 medium (5 mM NaCl, 0.17 mM KCl, 0.33 mM CaCl_2_·2H_2_O, and 0.33 mM MgSO_4_·7H_2_O). The embryos were staged according to the standard developmental staging guidelines (Kimmel et al., 1995).

### DNA constructs, RNA synthesis and injection

The transient expression constructs of *pCS2+-nphp3N-AKAR-CR*, *pCS2+-cPKA/dnPKA*, *pCS2+-nphp3N/(G2A)-cPKA/dnPKA-eGFP, pCS2+-cPKA/dnPKA-mRAB23(S23N/Q68L), pCS2+-IFT88-Flag* and the transgenic construct of *pMiniTol2-*β*-actin-npnp3N-AKAR-CR* and *pMiniTol2-en2a-eGFP* were generated by Gibson assembly. The primers used for cloning are listed in Table S2. The transient expression constructs of *pCS2+-SMO-eGFP*, *pSP64-shh* and *pDB600* were used as described (Maurya et al., 2013; Zhao et al., 2016).

The vectors *psPAX2* and *pMD2G* were kindly provided by the lab of Jianguo Li (Institute of Biomedical Sciences, Shanxi University, Taiyuan, China). cPKA was a corresponding mutation (H89Q; W198R) in the catalytic subunit of zebrafish PKA as previously described (Orellana et al., 1992). dnPKA was modified in accordance with previous study (Clegg et al., 1987).

For capped mRNA synthesis, the transient expression constructs were linearized by Not I for *pCS2+* constructs, Bam HI for *pSP64-shh* and XbaI for *pDB600* plasmid and then purified by Zymogen DNA concentrator. The mRNA was produced using either Sp6 or T3 Invitrogen Ambion mMessagemMachine® Kit according to the promoter in the constructs following the instructions, then purified mRNA was stocked in −80℃ for further use.

For transient expression of the fusion protein in zebrafish embryos, 150 pg of mRNA was injected into one-cell stage embryos.

### Generation of zebrafish transgenic lines

To generate the *sx1002* stable transgenic line, the *pMiniTol2-*β*-actin-npnp3N-AKAR2-CR* construct was co-injected with Tol2 transposase mRNA into one-cell stage embryos. T_0_ embryos with strong green and red fluorescence were screened at 24 hpf and inoculated as potential founders. Mature T_0_ fish were out-crossed with wild-type, and T_1_ offspring with consistent fluorescence were selected as stable transgenic lines.

To create the stable transgenic line Tg(*en2a:eGFP*)^sx1005^ (*sx1005*), Tol2 transposase mRNA was co-injected into one-cell stage embryos along with the *pMiniTol2-en2a-eGFP* construct (Maurya et al., 2013). T_0_ embryos with strong green-fluorescence were selected at 24 hpf and inoculated as potential founders. T_1_ embryos with steady fluorescence obtained by outcrossing of potential founders with wild type were selected as stable transgenic lines.

### FRET imaging

FRET assay was performed as described (Karpova et al., 2003). The NIH3T3 cells and the embryos were prepared. Fluorescence imaging was performed using the LSM 710 confocal inverted microscope (Zeiss). Time stacks of images (512 × 512 pixels, typical field of view 135 μm × 135 μm, 600 Hz scanning frequency) were acquired with a 63× oil immersion objective at room temperature under atmospheric conditions. The pinhole was set at 1AU. The Clover was excited at 488 nm, and its emission was detected in the range 500-550 nm and the mRuby emission was detected in the range >561 nm. Pre-experiments under the emission range of mRuby were carried out to test the bleaching effect. All ROIs are 90% bleached to maximize the bleaching of the acceptor with little or no effect on the donor. At least 2 regions were selected for each field, with one non-target set as the background and the random cell region set as the object. The acquisition of all channel images was set up as a time series of 5 cycles, with bleaching starting after 2 cycles.

For each sample, image time series have been acquired by selecting the field of view populated with more than 40 cells. After acquisition, data have been analyzed through the FRET package of Zen Imaging Software. The ROIs including each cell were selected and the Clover and mRuby fluorescence intensity at different time points before and after bleaching in both acquisition channels were calculated. The relative FRET efficiency was calculated with the following formula: FRET=100 × (IF _Donor, after_ - IF _Donor, before_) / IF _Donor, after_. The raw data were then further elaborated with Excel™.

### Whole-mount in situ hybridization

The probes of *ptch2*, *olig2*, and *nkx2.2a* for in situ hybridization were prepared as described (Barth and Wilson, 1995; Concordet et al., 1996; Stamataki et al., 2005), and *in situ* hybridization was performed as described (Thisse and Thisse, 2008). Images were captured using Zeiss Imager M2 microscope.

### Western blot analysis

Total protein of embryos and culture cells was prepared as described (Maurya et al., 2013; May et al., 2021). In brief, the embryos were lysed using lysis buffer (20 mM Tris HCl (pH 7.4), 150 mM NaCl, 1% TritonX-100, 10% Glycerol, 2 mM EDTA, and 1 mM PMSF) at the ratio of 1uL lysis buffer per embryo. The culture cells were lysed using RIPA buffer (50 mM Tris-HCl at pH 7.5, 150 mM NaCl, 1% Triton X-100, 1% sodium deoxycholate, 0.1% SDS, 1 mM EDTA, and 1X protease inhibitors). The supernatant was collected after centrifuged at 12, 000 g for 20 min at 4, and then was mixed with loading buffer (37.5 mM Tris HCl, pH 7.4; 3% SDS; 0.01% bromophenol blue; 6.25% glycerol; and 100 mM DTT) and boiled for 5 min before PAGE gel assay. Total protein of 40 embryos were separated on 10% acrylamide gel, and then the proteins were transferred onto the PVDF membrane (GE Healthcare Life Science, 10600023). The blocking and antibody incubation were performed accordingly. All antibodies used and their dilution are listed in Table S1. The secondary antibodies labeled by HRP were detected with the super sensitive ECL substrate kit (Thermo Fisher Scientific, 34580). And the images of the western blot were captured using an infrared imager (GE Healthcare).

### Immunofluorescence

Immunofluorescence (IF) staining on embryos was performed as described (Ben et al., 2011). In brief, the embryos were fixed with 4% paraformaldehyde (PFA) at room temperature for 2 h and then stored at −20°C with absolute methanol after dehydration. To proceed IF, the fixed embryos were permeabilized in acetone after rehydrated. Then the embryos were subsequently incubated by blocking buffer (DPBS, 1% BSA, 1% DMSO, 0.5% TritonX-100), primary and secondary antibodies in blocking buffer, and then washed in PBTX (DPBS, 0.5% TritonX-100). The washed embryos were stored in 70% Glycerol/DPBS at 4°C. The antibodies and their titration used in this study were listed in Table S1.

For cell IF staining, the cells were prepared on 14mm coverslips (NEST, 801010), and were fixed with 4% PFA. The IF staining was performed as described (Tsurumi et al., 2019), with the blocking buffer of 1% bovine serum albumin (BSA) in PBS, and antibodies in PBS. The antibodies and their titration used in this study were listed in Table S1. The cells on the coverslips were mounted onto microscope slides using the antifade reagent (Beyotime Biotechnology, P0131), after IF staining. The imaging was conducted using LSM 710 confocal microscope (Zeiss Microsystem, Germany).

## Statistics

All experiments were conducted with a minimum of three replicates. Results were displayed as the mean ± standard deviation. *P*-values were determined using one-way ANOVA with GraphPad Prism 7.0 software, and the difference with *P*-value less than 0.05 was set as the threshold for statistical significance. All image processing and analysis were performed in ImageJ (Version v1.52i, U.S. National Institutes of Health, Bethesda, Maryland, USA). Plots and Figures were generated using GraphPad Prism (Version 7.0, GraphPad Software, Inc.) and Adobe Illustrator CS5 (Version v15.0.0, Adobe Systems, Inc.).

## Ethical Statement

All animal experiments were conducted in accordance with the guidelines and regulations for the care and use of laboratory animals, and were approved by the Institutional Animal Care and Use Committee of Scientific Research in Shanxi University (CSRSX) (Approval No. [SXULL2019005]). Efforts were made to minimize animal suffering and to reduce the number of animals used, in alignment with the principles of the 3Rs (Replacement, Reduction, and Refinement).

## Acknowledgement

We are grateful to Michael Lin who kindly deposited the *AKAR2-CR* plasmid used for this study in Addgene, to Changxin Wu for providing the high-level experiment platform. This work was generously supported by National Natural Science Foundation of China (No. 31970738, U21A20389 and No. 81960286), Central Guidance on Local Science and Technology Development Fund of Shanxi Province, (YDZJSX2024D009).

## Authors’ contributions

H.Z.: data curation, investigation, methodology, project administration, software, validation, writing—original draft; Z.H., S.C. and J.B.: data curation, investigation, methodology, visualization; G.C.: conceptualization, methodology, writing—review and editing; P.W.I.: conceptualization, funding acquisition, resources, supervision, validation, writing—review and editing; Z.Z.: conceptualization, funding acquisition, methodology, project administration, resources, supervision, validation, visualization, writing—review and editing. All authors gave final approval for publication and agreed to be held accountable for the work performed therein.

## Supplemental Files

**Figure S1.**
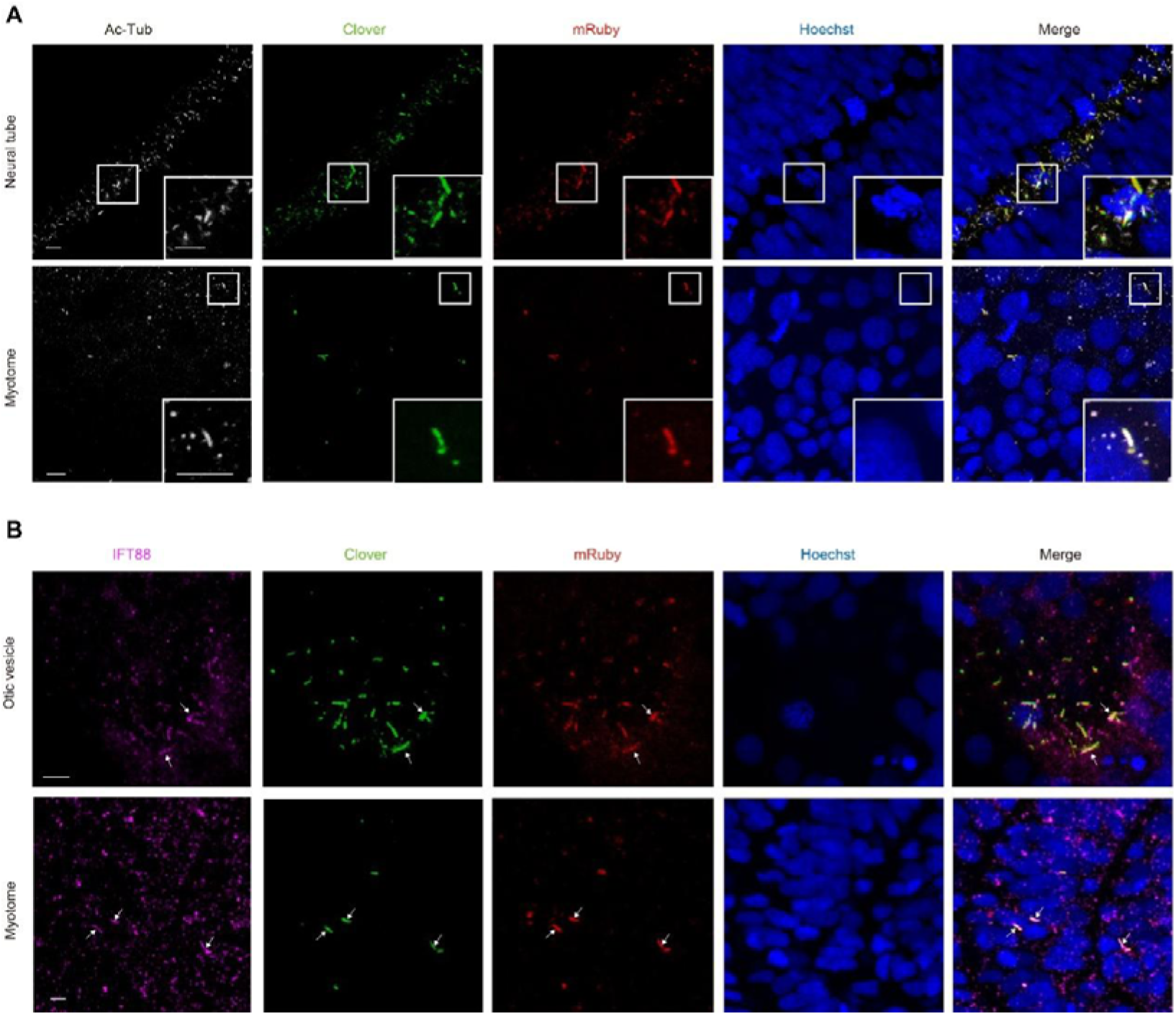
The expression of Nphp3N-AKAR2-CR does not affect the ciliary structure or intraflagellar transport in zebrafish. (A) The transiently expressed zNphp3N-AKAR2 in zebrafish embryos as indicated by Clover in green and mRuby in red was efficiently localized in the primary cilia of cells in the neural tube and myotome. Cilia were labeled by Ac-Tub in gray. Each small frame on the lower right corner of the main frame denotes the enlarged site as encircled by white square. The nuclei were labelled by Hoechst in blue. Scale bar, 5 μm. (B) Confocal images show that ciliary transport functions normally in the *sx1002* transgenic zebrafish line. The magenta channel indicates that IFT88 expression is detectable in both the otic vesicles and neural tubes of the transgenic zebrafish. In the *sx1002*, fluorescence signals are indicated by Clover in green and mRuby in red. The nuclei were labelled by Hoechst in blue. Arrows denote distinct IFT88 signals within the cilia of the transgenic line. Scale bar, 5 μm.

**Figure S2.**
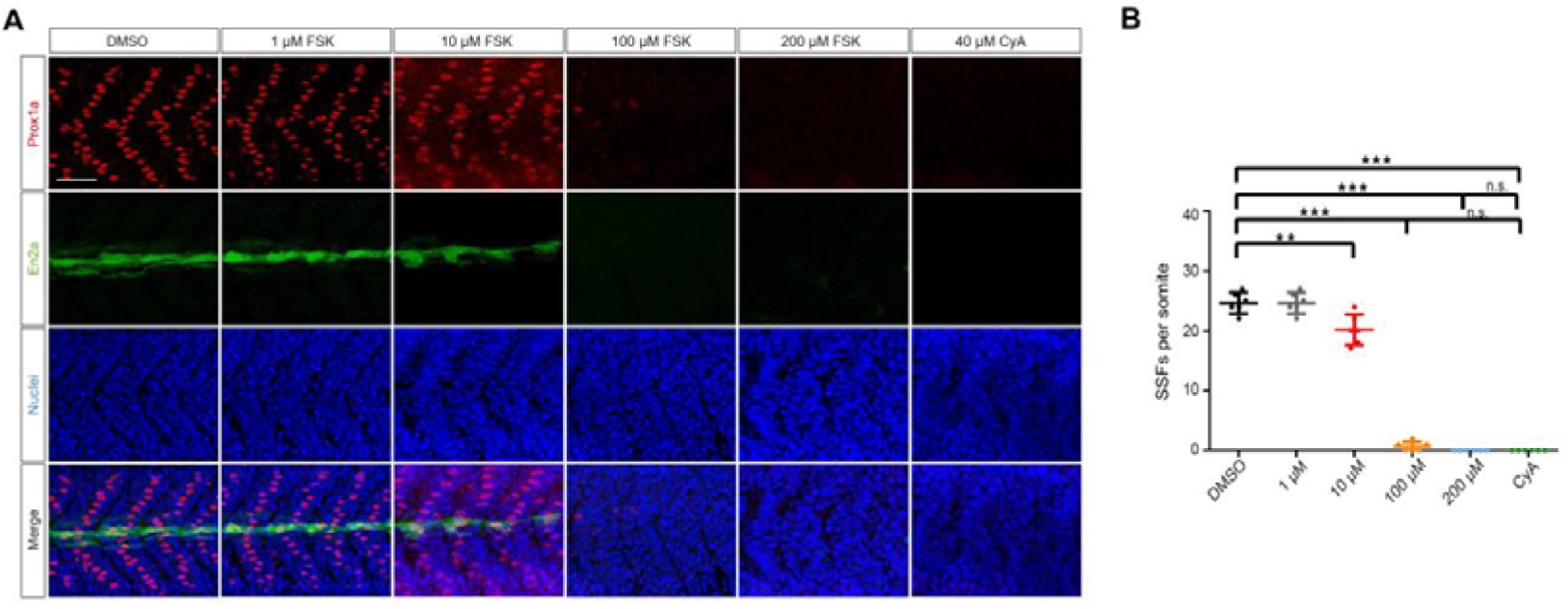
Treating zebrafish embryos with forskolin reduced the expression of Prox1a and En2a in the developing somite. (A) The activity of Hh pathway was indicated by expression of Prox1a in red and En2a:eGFP in green in the embryos of Tg (*en2a*:*eGFP*)^sx1005^, a reporter of Hedgehog signal transduction. Embryos were treated with increasing doses of FSK and 40 μM cyclopamine (CyA) from 6 to 24hpf, respectively. The nuclei were labelled by Hoechst in blue. Scale bar, 50 μm. (B) Quantification of Prox1a+ cells from experiments presented in A (*n* = 6 somites in 3 embryos). One-way ANOVA was used for analysis. ns indicates ‘not significant’ with *P*>0.05, and *** indicates ‘significant’ with *P*<0.001.

**Figure S3.**
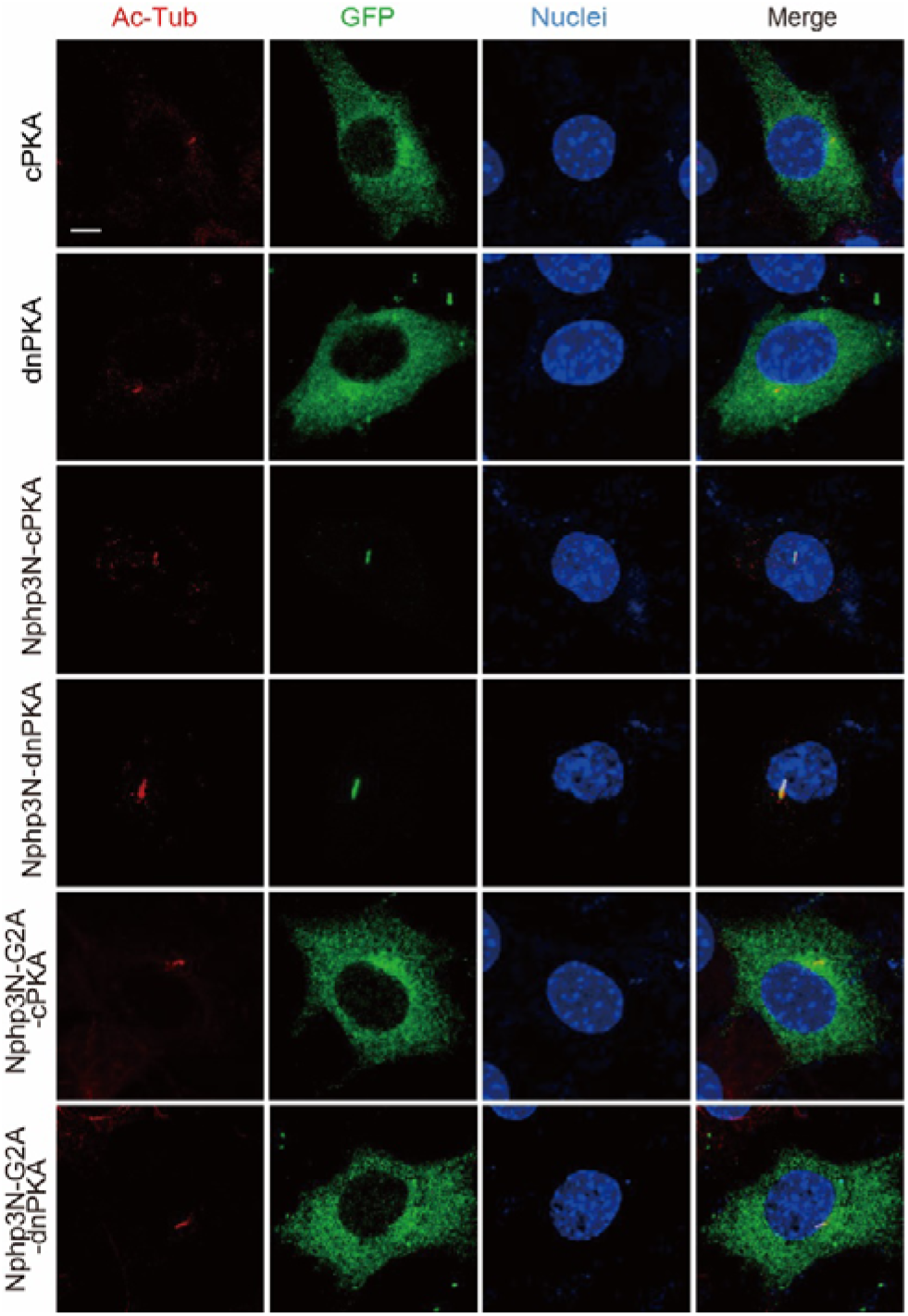
Nphp3N specifically drove cPKA/dnPKA to the PC in NIH3T3 cell. Subcellular localization of the different forms of cPKA/dnPKA when transiently expressing in NIH3T3 cells, as indicated by eGFP in green. The cilia were labeled by Ac-Tub in red and the nuclei by Hoechst in blue. Scale bars, 5 μm.

**Figure S4.**
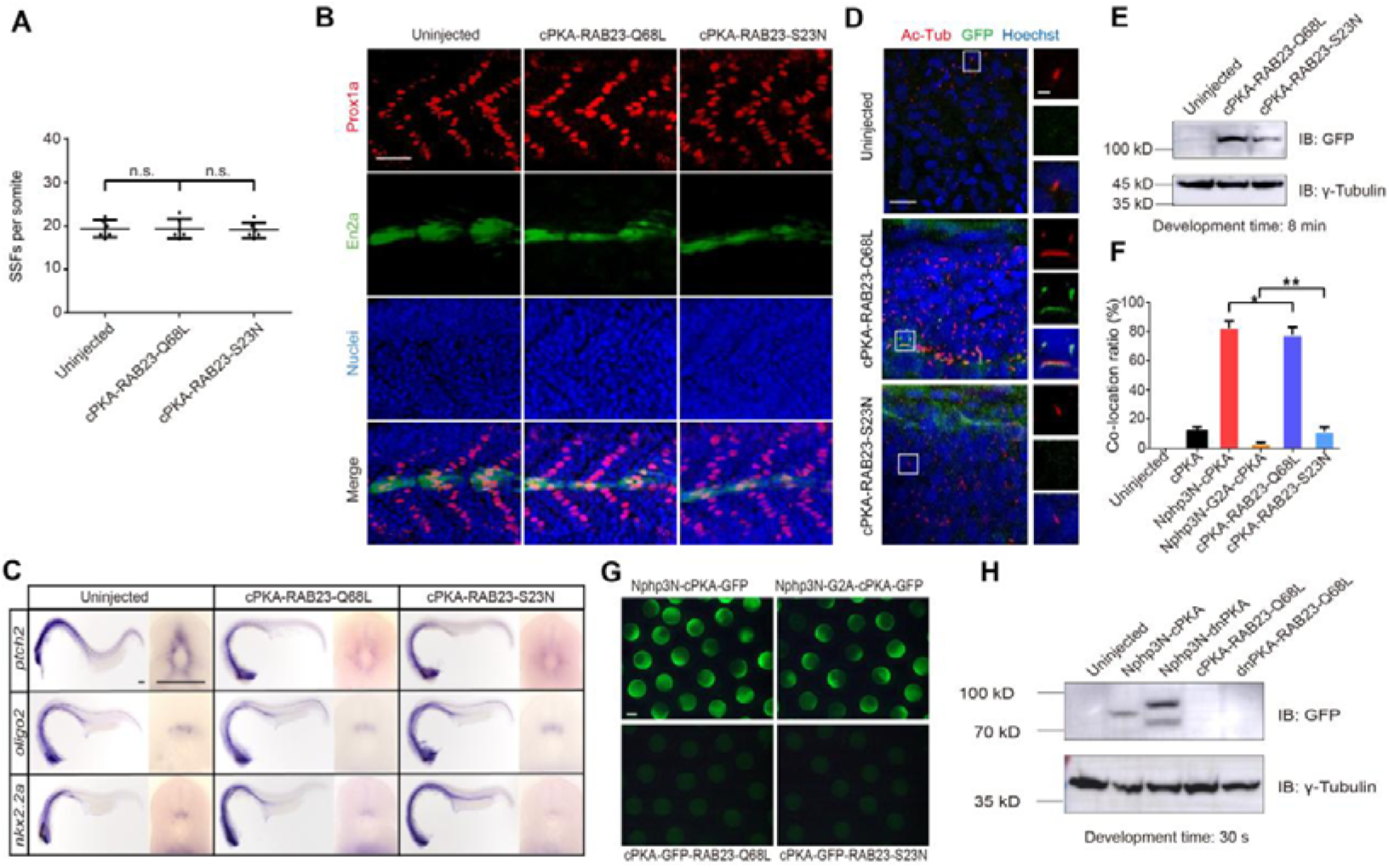
The cPKA driven by the RAB23 variant only weakly regulates the HH pathway. (A) Quantification of Prox1a+ cells from experiments presented in B (*n* = 6 somites from 3 embryos). One-way ANOVA was performed. And ns indicates ‘not significant’ with *P*>0.05.(B) The activity of Hh pathway was indicated by expression of Prox1a in red and En2a:eGFP in green in the embryos of Tg (*en2a*:*eGFP*)^sx1005^ expressing indicated RAB23 (Q68L/S23N)-cPKA. The nuclei were labelled by Hoechst in blue. Scale bar, 50 μm. (C) *In situ* hybridization of *ptch2*, *nkx2.2a* and *olig2* on the 24 hpf embryos expressing indicated cPKAs. Each panel showed a full view of the embryo on the left and a cross-sectional view of a somite on the right (*n* = 3 for each sample). Scale bars, 100 μm. (D) Subcellular localization of the cPKA-eGFP-Rab23 Q68L/S23N in embryos at 18 hpf, as indicated by eGFP in green. The cilia were labelled by Ac-Tub in red, and the nuclei by Hoechst in blue. Frames on the right panel indicates cilia depicted in the inset, with cilia on the top, indicated cPKAs/dnPKAs in the middle, and merge images at the bottom. Scale bars, 10 μm for the left panel and 2.5 μm for the right panel. (E) Immunoblot of lysates from 18 hpf zebrafish embryo expressing indicated GFP-tagged forms of RAB23-cPKAs. The γ-tubulin was used as loading control. Development time, 8 min. (F) Quantification of ciliary colocalization from experiments presented in Fig 2B and D (*n* = 60 cilia in 3 embryos). One-way ANOVA was used for analysis, and ** indicates ‘significant’ with *P*<0.01, * indicates ‘significant’ with *P*<0.05. (G) Transient expression of the indicated Nphp3N (WT/G2A)-cPKA and RAB23 (Q68L/S23N)-cPKA in zebrafish embryos at 6 hpf were indicated by eGFP in green. Scale bars, 500 μm. (H) Immunoblot of the indicated fusion proteins in embryos. The γ-tubulin was used as loading control. Development time, 30 s.

**Figure S5.**
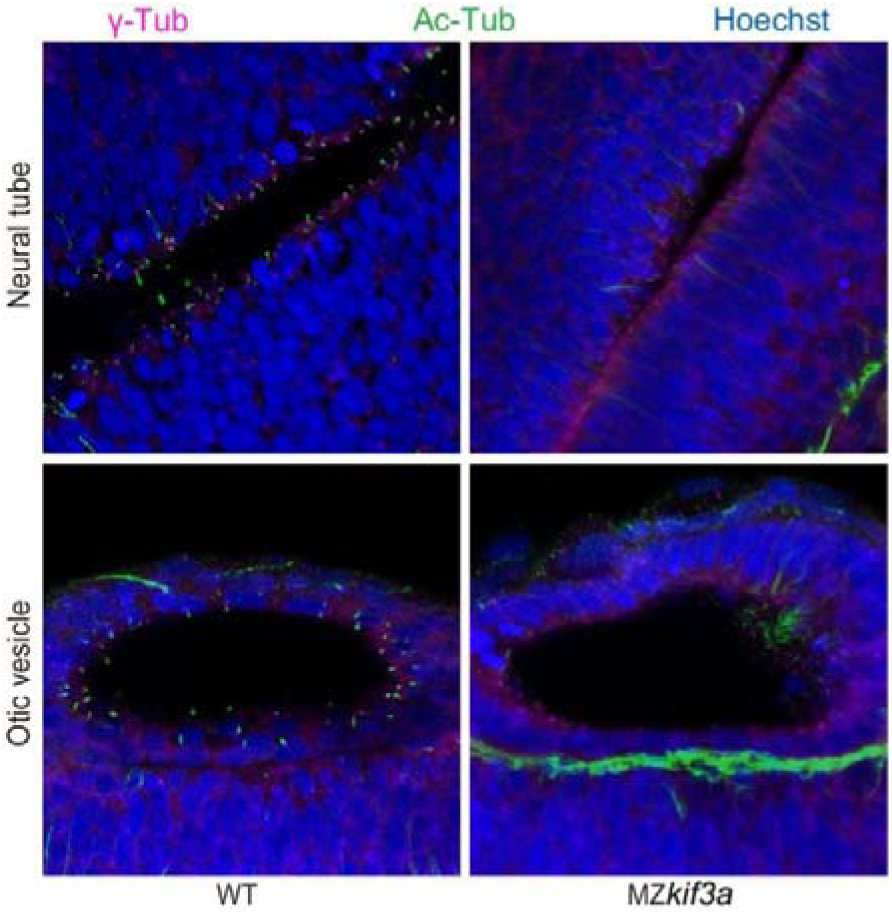
Confocal images showing cilia in the neural tube and otic vesicle of wild type and MZ*kif3a* mutants as indicated. Cilia were labelled with acetylated tubulin in green and the basal bodies were stained with gamma tubulin in magenta. Nuclei were stained with Hoechst in blue. Scale bar, 10 μm.

**Figure S6.**
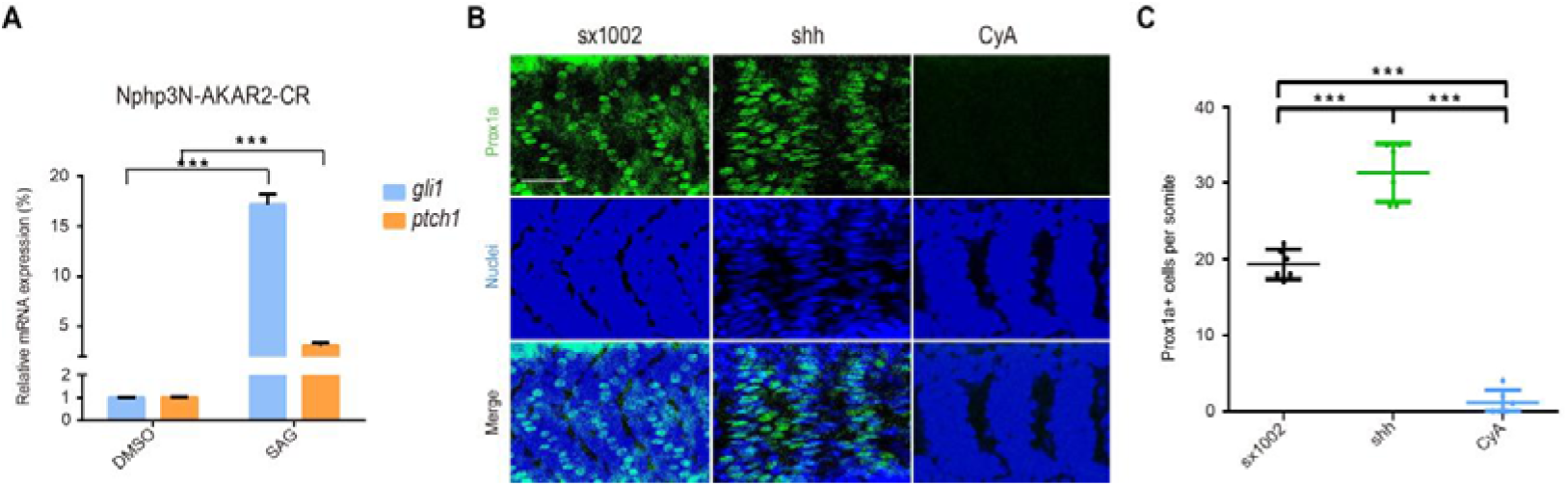
The expression of Nphp3N-AKAR2-CR does not affect the function of HH modulators. (A) NIH3T3 cells were transfected with Nphp3N-AKAR2-CR, then treated with 1 μM SAG or an equivalent volume of DMSO, respectively. Expression levels of *gli1* and *ptch1* were subsequently quantified. Data represents the mean and ± SD (n = 3). One-way ANOVA was used for analysis, and *** indicates ‘significant’ with P<0.001. (B) Representative Prox1a (red)/nuclei (blue) immunostainings of *sx1002* embryos at 24 hpf after injected with *shh* mRNA or treated with CyA, respectively. Scale bars, 50 μm. (C) Quantification of Prox1a+ cells from experiments presented in B (*n* = 6 somites from 3 embryos). One-way ANOVA was performed. And *** indicates significant with *P*<0.001.

